# Ketamine potentiates a central glutamatergic presynapse

**DOI:** 10.1101/2023.12.17.571741

**Authors:** Abdelmoneim Eshra, Noa Lipstein, Stefan Hallermann

**Affiliations:** Carl-Ludwig-Institute for Physiology, Medical Faculty, University of Leipzig, Liebigstrasse 27, 04103 Leipzig, Germany; Leibniz-Forschungsinstitut für Molekulare Pharmakologie (FMP), 13125 Berlin, Germany

## Abstract

Ketamine produces rapid and sustained antidepressant effects after brief exposure to a single dose. Counterintuitively, while ketamine acts primarily as a blocker of postsynaptic N-methyl-D-aspartate receptors (NMDARs), increased signalling at glutamatergic synapses has been reported. Due to technical limitations, however, it remains unclear whether ketamine directly increases presynaptic glutamate release or acts via postsynaptic or network-level mechanisms. To address this knowledge gap, we used presynaptic capacitance measurements to directly monitor glutamate release in a cerebellar synapse. Ketamine increased glutamate release within minutes and this effect persisted >30 minutes after washout. MK-801, another NMDAR blocker, had no effect on glutamate release. Mechanistically, we show that the ketamine-mediated enhancement of presynaptic release results from an increase in both calcium influx and the number of release-ready vesicles. Our data uncover a rapid effect of ketamine on key presynaptic properties of central glutamatergic synapses, which has important implications for the development of antidepressant drugs.

## Introduction

Ketamine has long been used in medicine to induce and maintain anaesthesia. In addition, recent studies in mice and humans have shown that ketamine has antidepressant effects at lower, sub-anaesthetic doses (1). For example, a single subanesthetic dose of ketamine infusion rapidly improved depressive symptoms in individuals with major depressive disorder, with antidepressant effects lasting three to seven days (2). The US Food and Drug Administration (FDA) approved intranasally administered ketamine (its S-enantiomer) to treat treatment-resistant depression in adults in 2019 and to treat major depressive disorder with acute suicidal intentions in adults in 2020 (3). The rapid and sustained antidepressant effect of ketamine may represent the biggest breakthrough in the field of psychiatry in the last 50 years (4).

Ketamine’s mode of action has been the subject of intense research, with several theories describing its potential mechanism of action (5, 6). There is general agreement that ketamine is an open channel blocker of NMDARs (7) and that the effects of ketamine involve neural plasticity, as the drug induces persistent changes in brain function despite a short elimination half-life (8). Ketamine induces a glutamate surge and a potentiation of glutamatergic signalling that results in an antidepressant effect (9–13) (but see evidence for decreased glutamatergic signaling (14)). However, the origin of the potentiation of glutamatergic synapses after ketamine application is still unclear.

A prominent theory for the potentiation of glutamatergic signalling is that ketamine disinhibits glutamatergic neurons through preferential blockade of NMDARs on GABAergic neurons (5, 6, 15, 16). This is sometimes called the “presynaptic effect” but refers to increased neuronal activity of glutamatergic neurons rather than to a direct effects of ketamine on presynaptic nerve terminals. The increase in net glutamate release is thought to induce postsynaptic potentiation of glutamatergic synapses, which has been extensively described at several synapses, both at the functional level (17–22) and at the structural level (16, 23, 24). Direct mechanistic effects of ketamine at presynaptic termainals have been proposed, but with controversial effects at different synapses and plasticity paradigms. For example, an effect of ketamine on presynaptic release probability, as assessed by the paired-pulse ratio, has yielded conflicting results at hippocampal synapses (25–27) and no change in release probability at prefrontal cortex synapses (28).

Our poor understanding of the presynaptic effects of ketamine is partly due to important technical limitations of the electrophysiological recording conditions: To our knowledge, all previous electrophysiological studies assessing the effects of ketamine on synaptic function have relied on postsynaptic measurements. In addition to the general limitations of relying on postsynaptic recordings to interpret presynaptic function, the case of ketamine is further complicated due to its function as an open NMDARs channel blocker (7) which could alter postsynaptic readouts. Moreover, the increased neuronal activity of glutamatergic neurons makes it difficult to disentangle direct effects of ketamine on glutamate release, intrinsic to the presynaptic terminal, from indirect effects on glutamate release due to increased network activity.

To clarify the role of ketamine in presynaptic function, we focused on the synapse between the mossy fibre and the granule cells of the mouse cerebellum. Although there is evidence that the cerebellum contributes to psychiatric disorders, including depression (29–31), we did not choose this synapse to investigate its role in the antidepressant effects of ketamine. Rather, this synapse is ideally suited for studying presynaptic glutamatergic transmission in the vertebrate brain in isolation: First, high-resolution presynaptic patch-clamp recordings and capacitance measurements are possible, allowing us to measure presynaptic release independent of postsynaptic glutamate receptors and independent of axonal mechanisms that may influence presynaptic action potentials (32–34). Second, the cell soma can be removed by cutting the axon during slice preparation, allowing us to isolate the immediate effects of ketamine on the presynapse, independent of upstream network effects acting on somato-dendritic compartments, and independent of major protein synthesis processes. Exploiting these technical possibilities, we show that presynaptic release is enhanced by both a profound increase in the number of release-ready vesicles and a small increase in presynaptic calcium influx soon after ketamine application, and that these effects are sustained long after ketamine washout.

## Results

### Ketamine rapidly enhances glutamate release

To investigate direct presynaptic effects of ketamine, we performed whole-cell patch clamp recordings from cerebellar mossy fiber boutons in parasagittal cerebellar slices in which the mossy fiber soma had been removed during preparation (Fig. 1a). To directly measure glutamate release independent of possible ketamine-induced postsynaptic changes, we recorded presynaptic membrane capacitance changes in response to depolarization pulses induced by the patch pipette. It has previously been shown at this synapse that the increase in presynaptic membrane capacitance is a linear measure of the amount of glutamate released (32) (Fig. 1b). We measured the change in presynaptic membrane capacitance upon applying depolarizing pulses of increasing duration from 1 to 100 ms in the presence of either control solution or 50 µM ketamine (Fig. 1c). The time in which the slice was incubated with ketamine prior to recording was approximately 30 min (Fig. 1d). During a whole-cell presynaptic recording, the release showed a low level of run-down (Supplementary Fig. S1). Comparison of the depolarization-induced capacitance increase in control and ketamine solution revealed a markedly increased glutamate release (Fig. 1e; see Supplementary Table S1 for p-values of two-way ANOVA and post-hoc Tukey tests). The mean value of the 10-ms-evoked capacitance jump normalised to resting capacitance (ΔC_m_/C_m_) was 70% greater in ketamine-treated nerve terminals compared to control nerve terminals (0.0289 for ketamine vs. 0.0169 for control; Fig. 1e). These data indicate that 50 µM ketamine strongly enhances glutamate release within 30 min.

**Figure 1.**
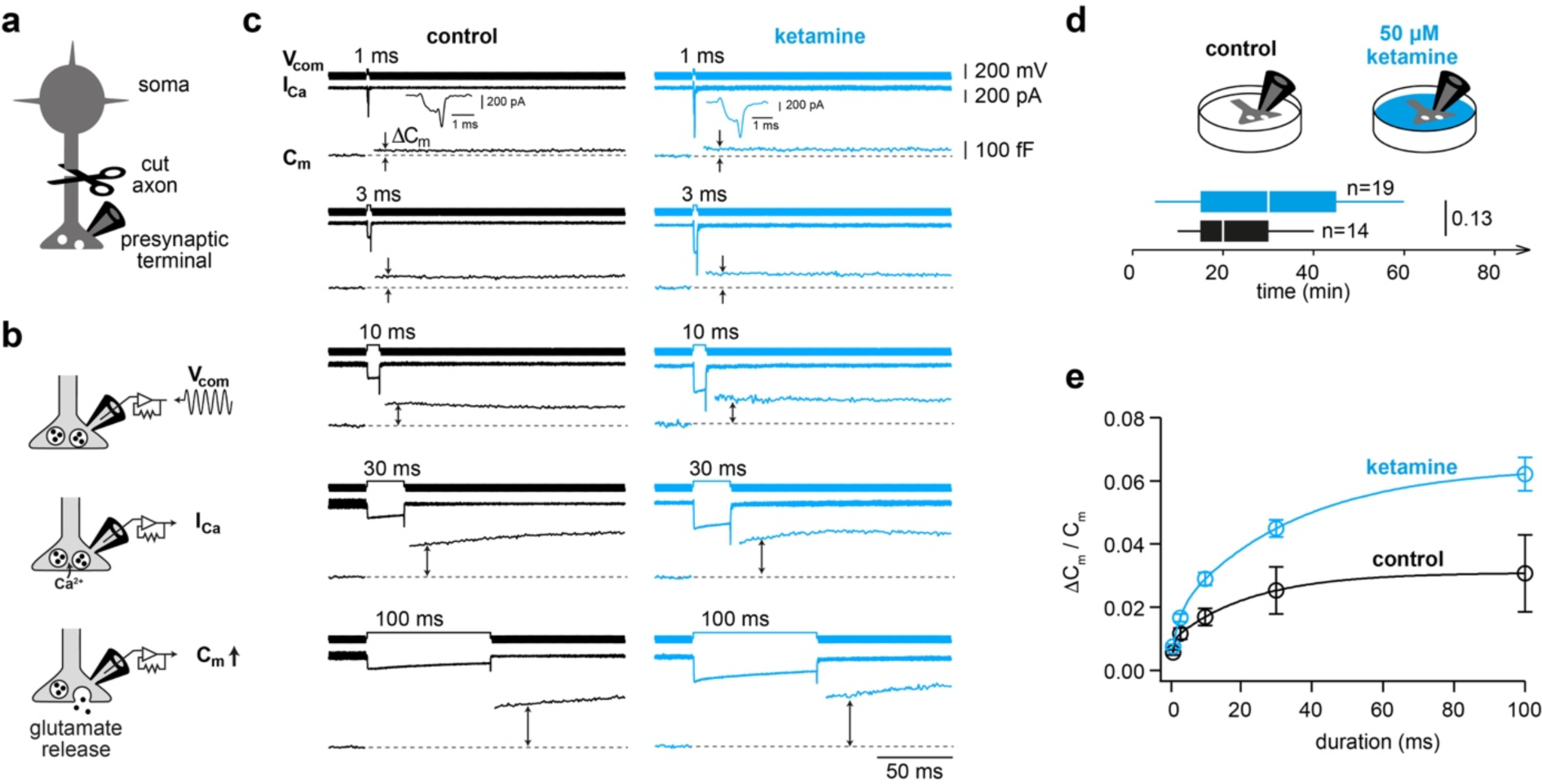
Ketamine rapidly enhances glutamate release. a. Pictogram of the recording configuration of the presynaptic terminal with a cut axon. b. Illustration of monitoring glutamate release with presynaptic capacitance measurements. *Top:* A sinusoidal voltage commend (V_com_) is applied. *Middle:* Depolarization induced presynaptic calcium current (I_Ca_) is measured. *Bottom:* Vesicular fusion and consequent glutamate release increases the presynaptic surface area and thus electrical membrane capacitance (C_m_). c. Example traces of the voltage commands used for applying the depolarizing pulses of different durations (between 1 and 100 ms) and measuring the pharmacologically isolated depolarization-evoked calcium current, and the depolarization-evoked capacitance jump for control (*left*) and ketamine application (*right*). d. *Top:* Illustration of acute application of ketamine. *Bottom:* Boxplots of the duration of ketamine application (median and interquartile range with whiskers indicating the whole data range). e. Average ΔC_m_ normalized to resting capacitance versus depolarization duration. Lines represent biexponential fits and the data are shown as mean with errors bars denoting standard error of the mean (SEM). Black and blue colors represent control and ketamine conditions, respectively. See Supplementary Table S1 for statistics.

### Sustained presynaptic potentiation after ketamine removal

To investigate whether the ketamine-induced presynaptic potentiation persists after ketamine removal, we performed another set of experiments in which brain slices were incubated for 30 min with either 50 µM ketamine or control solution. The slices were then transferred to a ketamine-free solution in which electrophysiology was performed after approximately 30 min of washout (Fig. 2a). Again, the change in presynaptic membrane capacitance was measured by applying depolarising pulses of increasing duration from 1 to 100 ms (Fig. 2b). The presynaptic release was also enhanced after washout of ketamine (Fig. 2c). The magnitude and kinetic properties of the enhanced release were strikingly similar to those observed in experiments performed in the presence of ketamine (see Fig. 1e). The mean value of the 10-ms-evoked ΔC_m_/C_m_ was 59% greater in ketamine-treated nerve terminals compared to control nerve terminals (0.0212 for ketamine vs. 0.0133 for control, Fig. 2c). These data indicate that the enhancement of glutamate release persists for at least 30 min after ketamine washout.

**Figure 2.**
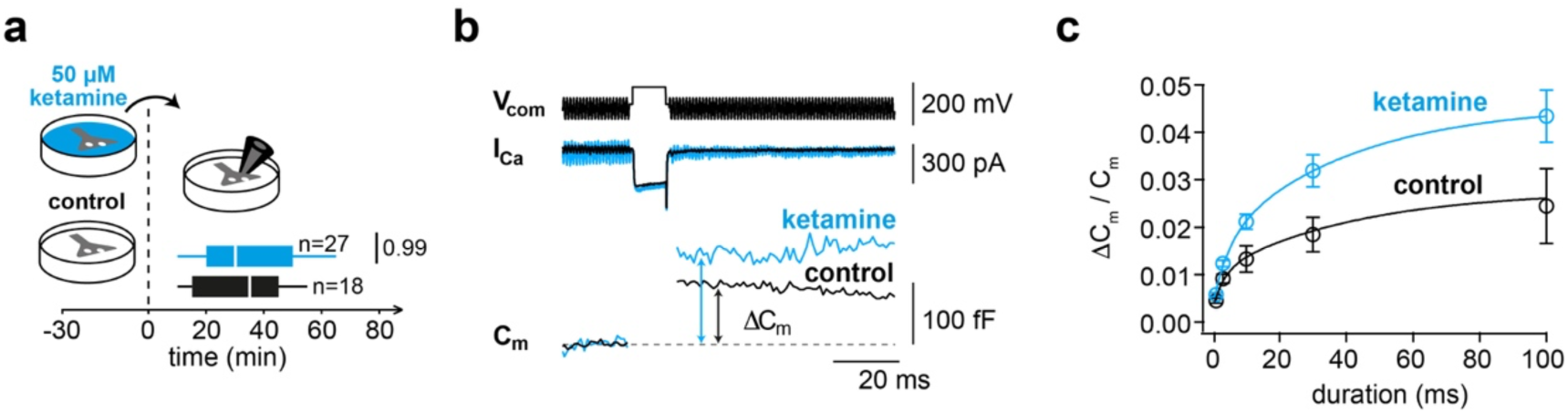
Sustained presynaptic potentiation after ketamine removal. a. Illustration of application of either 50 µM ketamine or control solution for 30 min and subsequent wash-out at time point t = 0 min. Boxplots show the duration of wash-out (median and interquartile range with whiskers indicating the whole data range). b. Example traces of the voltage command (V_com_), pharmacologically isolated calcium current (I_Ca_), and the capacitance jump (C_m_) for a duration of 10 ms for control (*black*) and ketamine (*blue*). c. Average ΔC_m_ normalized to resting capacitance versus depolarization duration. Lines represent biexponential fits and the data are shown as mean with errors bars denoting standard error of the mean (SEM). Black and blue colors represent control and ketamine conditions, respectively. See Supplementary Table S1 for statistics.

### MK801 does not increase glutamate release

To analyse whether NMDAR blockade can operate through a retrograde mechanism to potentiate presynaptic release, we used MK-801, an open channel NMDAR blocker like ketamine, which binds the channel at the same domain (PCP binding site) but with a longer trapping time. We incubated brain slices with either 50 µM MK-801 or control solution for 30 min, followed by approximately 30 min of washout (Fig. 3a). Monitoring glutamate release by presynaptic membrane capacitance measurements as described above, we observed that MK-801 had no effect on presynaptic glutamate release (Fig. 3b and c). To rule out that the lack of effect of MK-801 was not due to technical issues related to dissolution or potency of MK-801, we confirmed the blocking effect of the drug on isolated somatic NMDA currents measured in neuronal culture (Supplementary Fig. S2). These data indicate that the potentiation of glutamate release by ketamine is not caused by a simple blockade of NMDAR.

**Figure 3.**
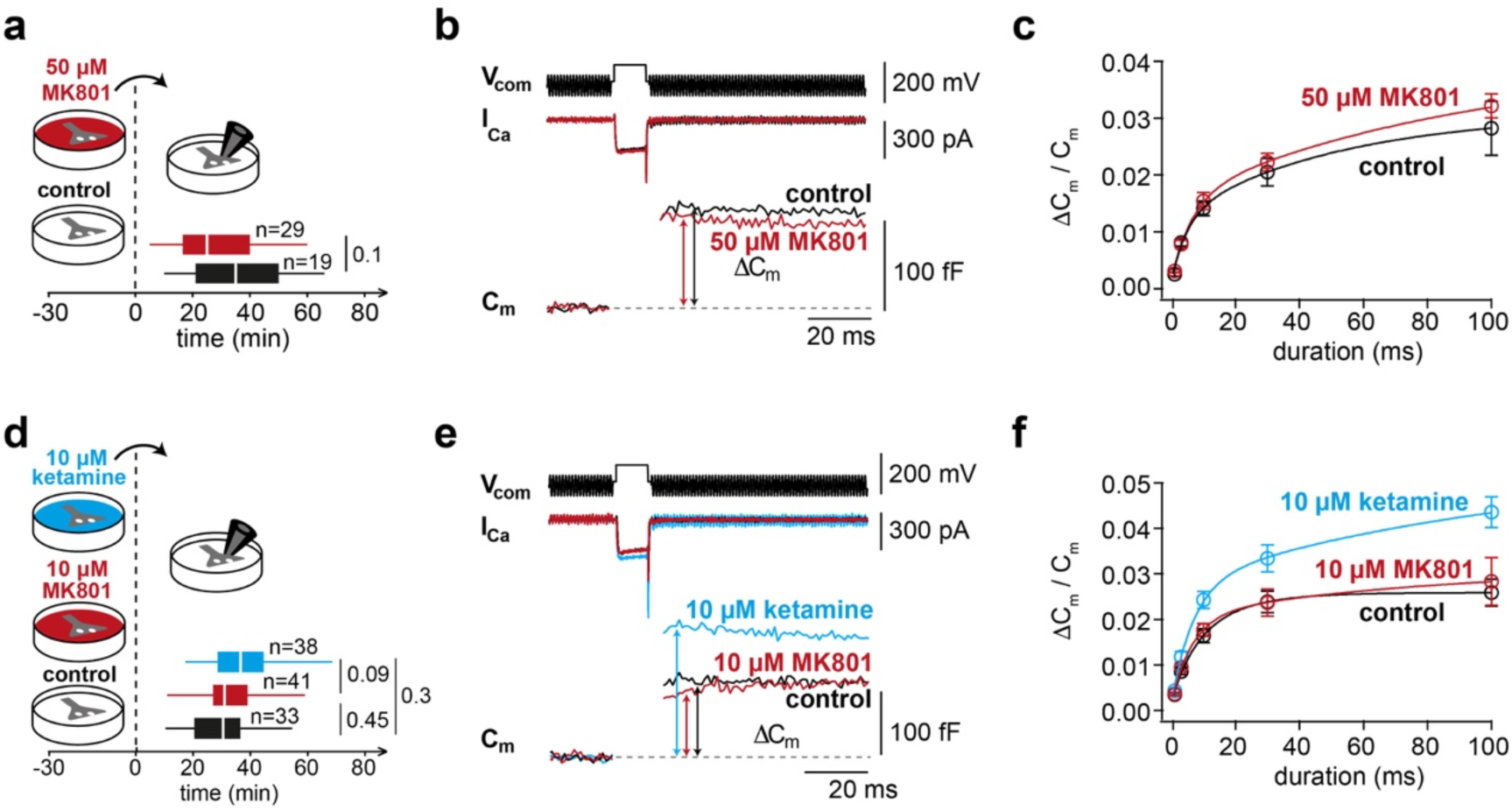
MK801 does not increase glutamate release. a. Illustration of application of either 50 µM MK801 or control solution for 30 min and subsequent wash-out at time point t = 0 min. Boxplots show the duration of wash-out (median and interquartile range with whiskers indicating the whole data range). b. Example traces of the voltage command (V_com_), pharmacologically isolated calcium current (I_Ca_), and the capacitance jump (C_m_) for a duration of 10 ms for control (*black*) and 50 µM MK801 (*red*). c. Average ΔC_m_ normalized to resting capacitance versus depolarization duration. Lines represent biexponential fits and the data are shown as mean with errors bars denoting standard error of the mean (SEM). Black and red colors represent control and 50 µM MK801 conditions, respectively. See Supplementary Table S1 for statistics. d. Illustration of application of either 10 µM ketamine, 10 µM MK801, or control solution for 30 min and subsequent wash-out at time point t = 0 min. Boxplots show the duration of wash-out (median and interquartile range with whiskers indicating the whole data range). e. Example traces of the voltage command (V_com_), pharmacologically isolated calcium current (I_Ca_), and the capacitance jump (C_m_) for a duration of 10 ms for control (*black*), 10 µ MK801 (*red*), and 10 µM ketamine (*blue*). f. Average ΔC_m_ normalized to resting capacitance versus depolarization duration. Lines represent biexponential fits and the data are shown as mean with errors bars denoting standard error of the mean (SEM). Black, red, and blue colors represent control, 10 µM MK801, and 10 µM ketamine conditions, respectively. See Supplementary Table S1 for statistics.

### Clinically relevant ketamine concentrations lead to presynaptic potentiation

The clinically-relevant ketamine concentrations in the brain are thought to be closer to 10 than 50 µM (20). We therefore conducted a set of experiments using 10 µM ketamine ketamine, 10 µM MK-801 or control solution applied for 30 min, followed by approximately 30 min of washout (Fig. 3d). After ∼30 min of washout, application of 10 µM ketamine increased depolarisation-induced presynaptic capacitance jumps by a similar amount compared to experiments with 50 µM ketamine (Fig. 3e and f). The mean value of the 10 ms evoked ΔC_m_/C_m_ was 48% greater in ketamine-treated nerve terminals compared to control nerve terminals (0.0243 for ketamine vs. 0.0164 for control, Fig. 3f). 10 µM MK-801 again did not show any presynaptic effects similar to the results using 50 µM MK-801.

To investigate how long the release-boosting effect of ketamine persists after washout of ketamine, we made use of the variable time needed to obtain the presynaptic recording configuration. There was no obvious decrease in the release-boosting effect of ketamine during the period of wash-out (Supplementary Fig. S3), indicating that the effect persists longer than 30 min.

### Ketamine enhances presynaptic calcium influx and calcium-to-release coupling

To explore potential mechanisms underlying the enhanced presynaptic release, we first analysed the steady-state amplitude of the pharmacologically isolated presynaptic calcium currents eliciting the capacitance jumps (Fig. 4a). In the three data sets, in which ketamine was used (Fig. 1, 2, and 3d-f), the calcium current density showed a consistent statistical trend towards higher amplitudes, although the effect did not reach significance in the individual data sets (Fig. 4b). When data from all experimental groups were pooled, regardless of concentration or method of drug administration, a significant increase in calcium current density was observed in ketamine-treated boutons compared to untreated boutons (Fig. 4b).

**Figure 4.**
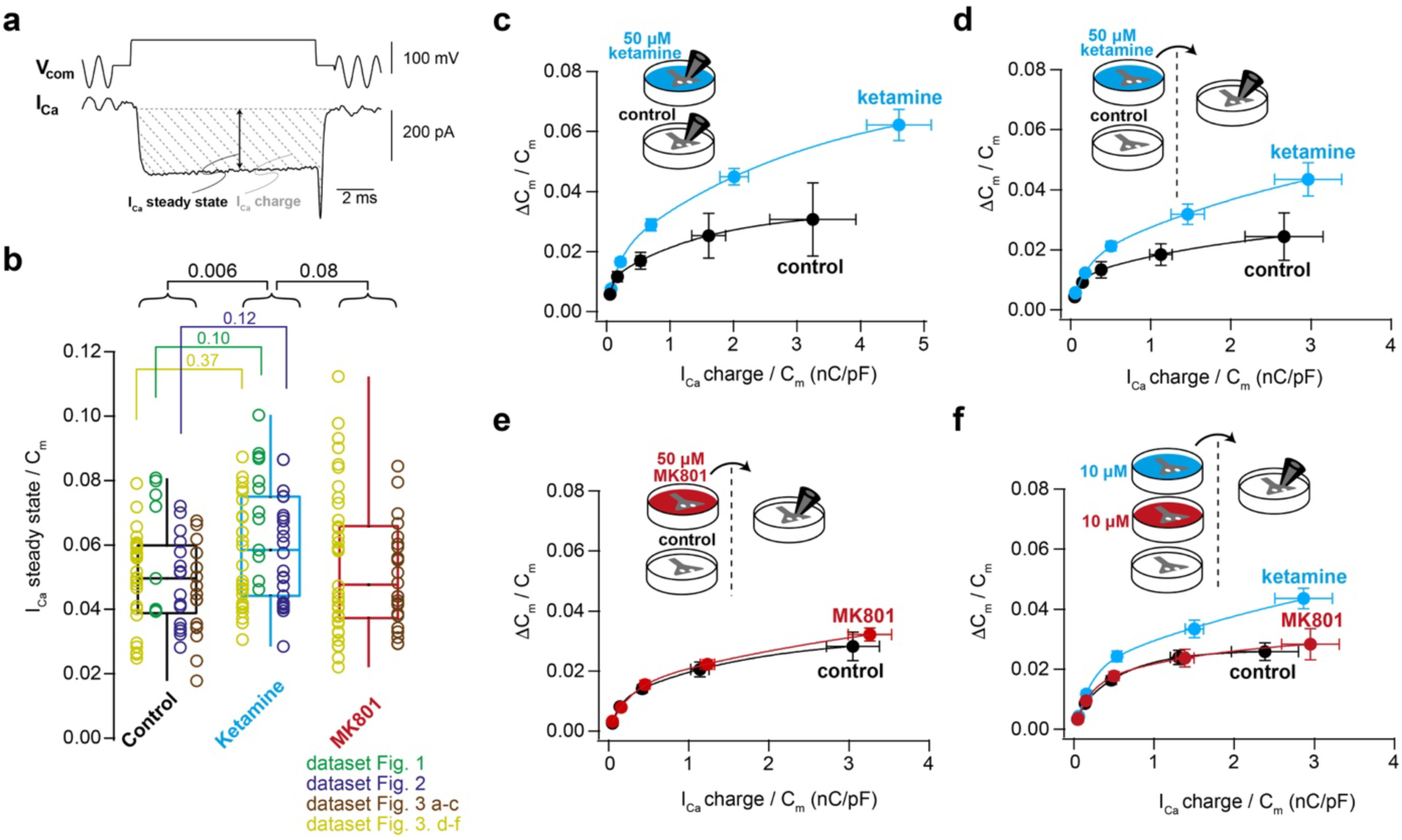
Ketamine enhances presynaptic calcium influx and calcium-to-release coupling. a. Example voltage command and calcium current evoked by a 10-ms depolarization. Dashed lines represent integral of the calcium current to measure the presynaptic calcium charge. The vertical arrow indicates the steady-state calcium current amplitude (measured in time windows of 2 ms before the tail currents). b. Merge of the steady-state amplitude of calcium currents evoked by a 3-ms depolarization from all experimental groups shown in Fig. 1-3. Color code represent different independent experimental groups. Boxplots show median and interquartile range with whiskers indicating the whole data range. Values of individual experiments are superimposed as circles. The numbers above the boxplots represent P-values as explained in the methods (see also Supplementary Table S1). c. Plot of average ΔC_m_ normalized to resting capacitance versus average charge transfer for experiments where the recordings were done in the presence of 50 μM ketamine. d. Plot of average ΔC_m_ normalized to resting capacitance versus average charge transfer for experiments where ketamine (50 μM) was applied for 30 with subsequent washout before the recording. e. Plot of average ΔC_m_ normalized to resting capacitance versus average charge transfer for experiments where MK-801 (50 μM) was applied for 30 min with subsequent washout before the recording. f. Plot of average ΔC_m_ versus average charge transfer for experiments where either ketamine 10 μM or MK-801 (10 μM) were applied for 30 min with subsequent washout before the recording. Lines in panel c-f represent biexponential fits and the data are shown as mean with errors bars denoting standard error of the mean (SEM). Black, blue, and red colors represent control, ketamine, or MK-801 conditions, respectively. See Supplementary Table S1 for statistics.

Given the release-calcium power relationship at this synapse (33), a slight increase in calcium influx could explain the enhanced release. We therefore analysed the relationship between release and calcium influx. For this purpose, we plotted ΔC_m_/C_m_ versus calcium charge, obtained by integrating the current over the entire pulse duration (Fig. 4a). These relationships were fitted with a biexponential function to describe the coupling between calcium and release (Fig. 4c-f). If the enhancement of presynaptic release was simply due to more calcium influx and hence more calcium charge, the ketamine data points should lie on the biexponential fit curve of the control data (i.e. more calcium influx would lead to more release within the normal calcium-release relationship at this synapse). In contrast, we found that ketamine, but not MK-801, resulted in a profound upward shift of the overall relationship (Fig. 4c-f), indicating that the efficiency of glutamate release upon calcium influx is increased by ketamine application. Therefore, our data suggest that ketamine increases presynaptic glutamate release both by increasing presynaptic calcium influx and by increasing the efficiency of calcium-release coupling.

## Discussion

We uncovered that ketamine directly potentiates presynaptic glutamate release in a rapid and sustained manner at a central synapse. The profound potentiation of glutamate release (∼70%) is independent of potential effects of ketamine on the cell body and on the neuronal network activity. Experiments with MK-801 indicate that the potentiation of release does not occur via simple NMDAR blockade. Our surprising finding of a profound ketamine-induced increase in glutamate release was based on four independent sets of technically challenging experiments comprising a total of 958 calcium-induced capacitance jumps from 238 presynaptic patch-clamp recordings (Fig. 1-3). Thus, our results provide robust evidence for a direct, rapid, and profound increase in presynaptic glutamate release at a central synapse.

The increase in glutamate release was sustained for >30 min after the wash-out of ketamine (Fig. 2, 3, and S3) indicating that ketamine induces a presynaptic form of plasticity (35). Although the time scale of 30 min to 1 h in our study is much shorter than the time scale of days for anti-depressant effect in patients, the here-observed enhanced presynaptic release can provide an explanation for the glutamate surge observed in patients and the ketamine induces neuronal-circuit potentiation (8–11, 36). The elevated glutamate release is thought to mediate the antidepressant effect of ketamine (8, 9) but further studies are needed to investigate the relationship between presynaptic potentiation and the antidepressant effect of ketamine.

Our data indicate that ketamine increases the number of release-ready vesicles rather than the release probability, because an increase in release probability should only potentiate release at depolarizations with short duration. Instead, ketamine potentiated release at all depolarization durations (1-100 ms). Our results are therefore consistent with studies showing no efffect of ketamine on paired-pulse ratio of postsynaptic currents (27). This often-used assessment of presynaptic strength is insensitive to changes in the number of release-ready vesicles. An increase in the number of release-ready vesicles is furthermore consistent with alterations of presynaptic vesicular proteins upon ketamine application (5, 37, 38), but this could also be due to indirect effects of elevated neuronal network activity.

Our finding that only ketamine but not MK-801 potentiates presynaptic function argues for an NMDAR-independent mechanism. Consistently, ketamine and MK-801 can differentially modulate cAMP and BDNF signalling (39). However, a role of NMDARs in the here-observed presynaptic potentiation cannot be fully excluded. Ketamine and MK-801 have different trapping time at the NMDAR, which might induce different current time courses or metabotropic-like effects (40). At the cerebellar mossy fiber synapse, a similar increase in glutamate release was recently observed upon homeostatic plasticity via blockade of postsynaptic AMPA receptors (41). Our MK-801 data argue against simple homeostatic plasticity as the cause of ketamine’s presynaptic potentiation, but more complicated forms of presynaptic homeostatic plasticity that depend on the used drug to block postsynaptic receptors (42) could mediate the presynaptic potentiation.

Interestingly, at CA3 neurons, ketamine has been shown to enhance the release probability in a HCN1 channel-dependent manner (26). Our results cannot be explained by a modulatory effect of ketamine on HCN1 function because the here-used voltage-clamp mode excludes effects of HCN channels on excitability (34). Furthermore, the metabolic product of ketamine, hydroxynorketamine (HNK), has been shown to mediate presynaptic strengthening (43). However, our results can also not be explained by the action of HNK because we applied ketamine directly to the brain tissue therefore excluding the metabolic pathway through the liver. Therefore, the here-described direct presynaptic potentiation is independent of postsynaptic signal amplification via NMDAR-blockade, circuit potentiation, NMDAR-dependent disinhibition, or ketamine metabolism, but could operate synergistically with these indirect mechanisms (5, 6, 18).

In conclusion, our data show that ketamine rapidly potentiates presynaptic function, therefore providing a novel mechanism for the ketamine-induced glutamate surge in patients. These results open up new avenues of research to address the urgent need for side effect-free rapid-acting antidepressants.

## Materials and Methods

### Sex as a biological variable

Our study examined male and female animals, and similar findings are reported for both sexes.

### Preparation

Animals were handled in accordance with European (EU Directive 2010/63/ EU, Annex IV for animal experiments), national and Leipzig University guidelines. All experiments were approved in advance by the federal Saxonian Animal Welfare Committee (T29/19). Acute cerebellar slices were prepared from mature P35–P42 C57BL/6 mice of either sex as previously described (33). Animals were anesthetized by isoflurane using a Dräger Vapor 2000 vaporizer (Dräger, Germany) and then sacrificed by decapitation. The cerebellar vermis was quickly dissected and mounted in a chamber filled with ice-cold artificial cerebrospinal fluid (ACSF). Parasagittal slices having a thickness of 300 µm were cut using a Leica VT1200 microtome (Leica Microsystems), transferred to an incubation chamber at 35 °C for ∼30 min, and then stored at room temperature until experiment time. The ACSF solution for slice cutting and storage contained (in mM) the following: NaCl 125, NaHCO_3_ 25, glucose 20, KCl 2.5, 2, NaH_2_PO_4_ 1.25, MgCl_2_ 1 (310 mOsm, pH 7.3 when equilibrated with Carbogen [5% (vol/vol) O_2_/95% (vol/vol) CO_2_]).

### Presynaptic recordings

Recordings were performed at physiological temperature as described (33). Borosilicate glass (2.0/1.0 mm outer/inner diameter; Science Products) was used to pull presynaptic patch-pipettes to an open-tip resistance of 3-5 MΩ (when filled with intracellular solution) using a DMZ Puller (Zeitz-Instruments, Germany). During recording, slices were superfused with ACSF containing (in mM): NaCl 105, NaHCO_3_ 25, glucose 25, TEA 20, 4-AP 5, KCl 2.5, CaCl_2_ 2, NaH_2_PO4 1.25, MgCl_2_ 1, and tetrodotoxin (TTX) 0.001, equilibrated with 95% O_2_ and 5% CO_2_. Cerebellar mossy fiber boutons were visualized with oblique illumination and infrared optics. Whole-cell patch-clamp recordings of cerebellar mossy fiber boutons were performed using a HEKA EPC10/2 amplifier controlled by Patchmaster software (HEKA Elektronik, Germany). All recordings were restricted to lobules IV–V of the cerebellar vermis to reduce potential functional heterogeneity among different lobules (55). The intracellular solution contained (in mM): CsCl 130, MgCl_2_ 0.5, TEA-Cl 20, HEPES 20, Na_2_ATP 5, NaGTP 0.3, ATTO-594 0.01.

Capacitance measurements were performed at a holding potential of −100 mV with sine-wave stimulation (5 kHz and ±50 mV amplitude) (32). During the sine wave stimulation, a brief depolarizing pulse was applied. Using an automated protocol written in Patchmaster, consecutive stimulations were performed and the duration of the depolarizing pulse in each stimulation was increased gradually (1, 1, 3, 10, 30, 100 ms). The duration between consecutive capacitance measurements was 15 s for the weak pulses (1 and 3 ms) and 30 s between the stronger pulses (10, 30, and 100 ms). To make sure that the waiting time between pulses is sufficient and that consecutive stimulation did not induce some sort of facilitation, we compared two consecutive 1 ms pulses at the beginning of each recording and found no difference (Supplementary Fig. 1). The hydrostatic pressure in the patch pipette pressure was kept at zero mbar.

### Drug application

For incubation experiments (Fig. 1b-d), brain slices were incubated with the drug of interest (ketamine or MK-801) at room temperature. After 30 min, the brain slice was transferred to a drug-free solution for a first washout step of about 10 s. Then, the slice was transferred to the recording chamber to perform electrophysiology while being continuously perfused with ketamine-free (and MK-801-free) extracellular solution. For experiments where the drug was continuously applied during recording (Fig. 1a), 50 μM ketamine was all the time included in the extracellular solution used for recording.

### Data analysis

All experiments and analyses (except for Supplementary Fig. 2) were performed in an interleaved and blinded manner. To this end, for each data set the aliquots of ketamine, MK-801, or ACSF were randomly, sequentially numbered by a second person and subsequentally added by the investigator. Unblinding was done after finishing the analyses. Analyses of capacitance measurements and calcium currents were performed using Patchmaster. To quantify the amplitude of capacitance jumps, we measured the difference between a baseline resting capacitance and the depolarization-evoked capacitance jump. The baseline capacitance was measured as the average capacitance during a time frame of 20-60 ms before the depolarizing pulse. the depolarization-evoked capacitance was quantified as the average capacitance during a time frame of 50-100 ms measured 20-30 ms after the end of the depolarizing pulse. Similarly, calcium current amplitudes were quantified by measuring the difference between a baseline (measured as the average current during a time frame of 2-20 ms before the depolarizing pulse) and a steady state current (measured as the average current during a time frame of 0.3-3 ms measured 0.5-3 ms after the end of the depolarizing pulse). Calcium charge transfer was obtained by integrating the current over the entire depolarizing-pulse duration.

### Statistics

For statistical comparisons of the data in Fig. 1e, 2c, 3c, 3f, and 4c-f, we perform two-way ANOVA and Tukey post hoc tests. For the data in Fig. 4b, a non-parametric one-way ANOVA (Kruskal–Wallis) test with non-parametric post hoc tests (Dwass–Steel– Critchlow–Fligner pairwise comparisons; DSCF) were used, but identical statistical significance levels were obtained with a parametric one-way ANOVA test and Tukey post hoc tests. For comparison of the individual data sets in Fig. 4b, Mann-Whitney U tests were used (color-coded P values are provided in the figure). The calculations were performed with jamovi (www.jamovi.org). See Table S1 for P-values.

## Author contributions

AE: project conceptualization, conducting experiments, analyzing data, writing the manuscript. NL: writing the manuscript. SH: project supervision, writing the manuscript.

## Acknowledgments

We would like to thank Stephen M. Smith, Volker Haucke, Christian Geis, and Jana Nerlich for critically reading the manuscript and Claudia Binder for technical assistance. This work was supported by the European Research Council (ERC Consolidator Grant 865634) to S.H. and by the German Research Foundation (DFG; FOR 3004 SYNABS, HA6386/10-2 to S.H. and Excellence Strategy EXC-2049-390688087 and CRC 1286 “Quantitative Synaptology” to N.L.).

## Figures and legends

**Supplementary Fig. S1.**
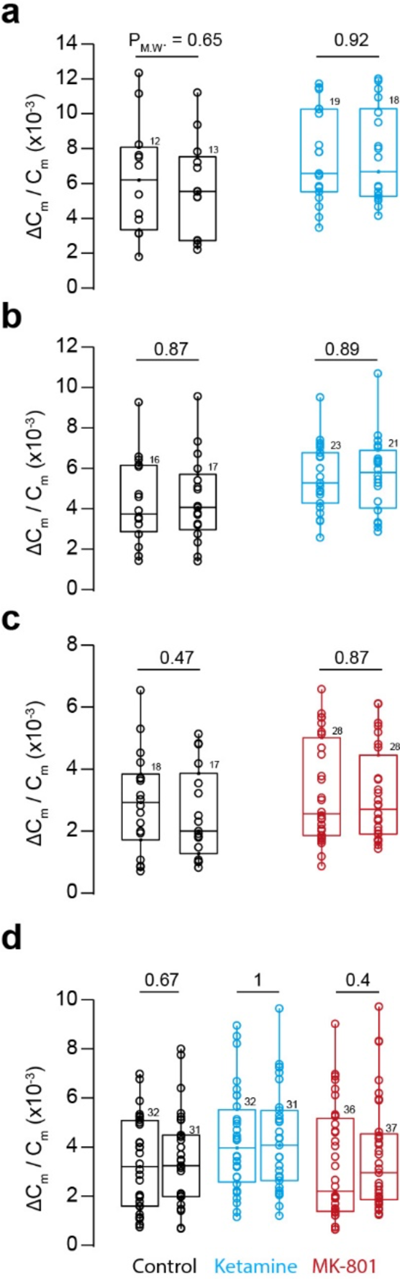
Amplitudes of ΔC_m_ do not differ between two consecutive 1 ms depolarisations. a. Boxplot comparison of two consecutive 1 ms pulses in experiments shown in Fig. 1a. left boxplots represent the first 1-ms-pulses while right boxplots represent the second 1-ms-pulses. b. Same comparison as in panel a but for experiments shown in Fig. 1b c. Same comparison as in panel a but for experiments shown in Fig. 1c d. Same comparison as in panel a but for experiments shown in Fig. 1d Black, red, and blue color code represent control, MK801, or ketamine conditions, respectively. Boxplots show median and interquartile range with whiskers indicating the whole data range. Values of individual experiments are superimposed as circles and the numbers of presynaptic recordings are written above each boxplot. The numbers above the boxplot-comparisons represent p-values of Mann-Whitney *U* tests.

**Supplementary Fig. S2.**
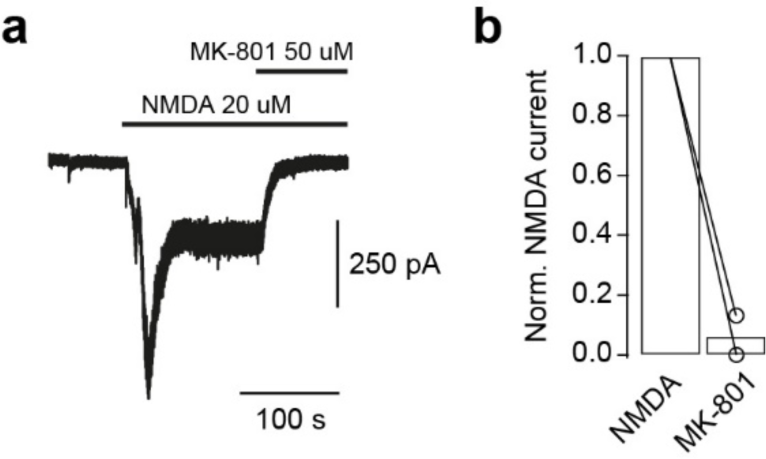
MK-801 blocks somatic NMDA currents in primary neuronal culture. a. Representative trace of a pharmacologically isolated inward current, induced via washing-in NMDA (20 μM), and blocked by washing-in MK-801 (50 μM). Currents were recorded in the presence of TTX (1 μM), NBQX (10 μM), glycine (3 μM), gabazine (10 μM), strychnine (200 nM). b. Comparison of the amplitude of the inward current (steady state) before and after applying MK-801 normalized to the NMDA-induced current (steady state).

**Supplementary Fig. S3.**
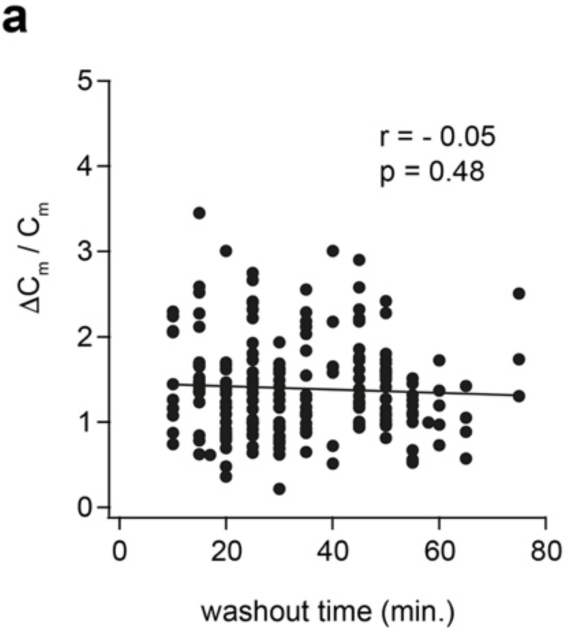
No dependence of release enhancement on washout time. a. Plot of ΔC_m_ /C_m_-values after ketamine incubation normalized to the average control value of the corresponding pulse plotted versus washout time. Merged data obtained from all stimulation pulses in Fig. 1b and d. r and p values are from Pearson correlation analysis.

**Supplementary Table S1.** Number of boutons recorded per data set and statistics. Only recordings where both calcium current and capacitance measurements could be analyzed were included. In cases where, the signal to noise ratio was not sufficient, clamping issues occurred, or the seal became leaky, the recording of the corresponding depolarization pulse was excluded. Therefore, the number of recordings varied among different depolarization pulses applied to each recorded bouton.

| Fig. | Drug | Number of recordings |  |  |  |  |  | P-value<br>two-way<br>ANOVA | P-value<br>Tukey<br>post hoc<br>test vs.<br>control |
| --- | --- | --- | --- | --- | --- | --- | --- | --- | --- |
|  |  | all | 1ms | 3ms | 10ms | 30ms | 100ms |  |  |
| 1e | Control | 14 | 13 | 9 | 8 | 6 | 4 | <0.001 |  |
|  | Ketamine | 19 | 18 | 12 | 10 | 8 | 8 |  | <0.001 |
| 2c | Control | 18 | 16 | 15 | 8 | 7 | 6 | <0.001 |  |
|  | Ketamine | 27 | 21 | 24 | 17 | 11 | 11 |  | <0.001 |
| 3c | Control | 19 | 17 | 16 | 13 | 11 | 11 | 0.15 |  |
|  | MK-801 | 29 | 28 | 24 | 19 | 17 | 17 |  | 0.15 |
| 3f | Control | 33 | 31 | 28 | 19 | 13 | 10 | <0.001 |  |
|  | Ketamine | 38 | 31 | 27 | 20 | 16 | 16 |  | <0.001 |
|  | MK-801 | 41 | 37 | 34 | 25 | 18 | 13 |  | 0.73 |
| 4c | Control | 14 | 13 | 9 | 8 | 6 | 4 |  |  |
|  | Ketamine | 19 | 18 | 12 | 10 | 8 | 8 |  |  |
| 4d | Control | 18 | 16 | 15 | 8 | 7 | 6 |  |  |
|  | Ketamine | 27 | 21 | 24 | 17 | 11 | 11 |  |  |
| 4e | Control | 19 | 17 | 16 | 13 | 11 | 11 |  |  |
|  | MK-801 | 29 | 28 | 24 | 19 | 17 | 17 |  |  |
| 4f | Control | 33 | 31 | 28 | 19 | 13 | 10 |  |  |
|  | Ketamine | 38 | 31 | 27 | 20 | 16 | 16 |  |  |
|  | MK-801 | 41 | 37 | 34 | 25 | 18 | 13 |  |  |
| Fig. | Drug | Number<br>of recor-<br>dings | P-value<br>Kruskal<br>-Wallis<br>test | P-value<br>DSCF<br>post hoc<br>test vs.<br>ketamine | P-value<br>one-way<br>ANOVA | P-value<br>Tukey<br>post hoc<br>test vs.<br>ketamine |  |  |  |
| 4b | Control | 68 | 0.007 | 0.006 | 0.006 | 0.004 |  |  |  |
|  | Ketamine | 63 |  |  |  |  |  |  |  |
|  | MK-801 | 58 |  | 0.08 |  | 0.14 |  |  |  |

